# Bayesian Modelling Approaches for Breath-Hold Induced Cerebrovascular Reactivity

**DOI:** 10.1101/2024.02.06.579134

**Authors:** Genevieve Hayes, Daniel P. Bulte, Stefano Moia, Martin Craig, Michael Chappell, Eneko Uruñuela, Sierra Sparks, César Caballero-Gaudes, Joana Pinto

## Abstract

Cerebrovascular reactivity (CVR) reflects the ability of blood vessels to dilate and constrict in response to a vasoactive stimulus and is an important indicator of cerebrovascular health. CVR can be mapped non-invasively with functional magnetic resonance imaging (fMRI) based on blood oxygen level-dependent (BOLD) contrast in combination with a breath-hold (BH) task. There are several ways to analyse this type of data and retrieve individual CVR amplitude and timing information. The most common approach involves employing a time-shifted general linear model with the measured end-tidal carbon dioxide signal as a regressor of interest. In this work, we introduce a novel method for CVR mapping based on a variational Bayesian approach. We analysed BOLD fMRI data from six participants that performed a BH task in ten different sessions each, and computed the corresponding CVR amplitude and delay maps for each session/subject. No statistically significant differences were observed between the modelling approaches in the CVR delay and amplitude maps in grey matter. Notably, the largest difference between methods was apparent in the case of low CVR amplitude, attributed to how each method addressed noisy voxels, particularly in white matter and cerebral spinal fluid. Both approaches showed highly reproducible CVR amplitude maps where between-subject variability was significantly larger than between-session variability. Furthermore, our results illustrated that the Bayesian approach is more computationally efficient, and future implementations could incorporate more complex noise models, non-linear fitting, and physiologically meaningful information into the model in the form of priors. This work demonstrates the utility of variational Bayesian modelling for CVR mapping and highlights its potential for characterising BOLD fMRI dynamics in the study of cerebrovascular health and its application to clinical settings.

## 1 Introduction

The maintenance of appropriate cerebral blood flow (CBF) is critical for brain function and survival, and is closely regulated by the contraction and dilation of cerebral blood vessels. Cerebrovascular reactivity (CVR) reflects this intrinsic mechanism of blood vessels to alter their calibre in response to vasoactive stimuli. Notably, CVR has been identified as an imaging biomarker of cerebrovascular health in a number of diseases including stroke (Krainik et al., 2005; Markus & Cullinane, 2001), brain tumours (Fierstra et al., 2018; Pillai & Zacá, 2011), carotid occlusion (Chang et al., 2009; Donahue et al., 2009; Webster et al., 1995), Alzheimer’s disease (Cantin et al., 2011; Glodzik et al., 2013; Suri et al., 2015), Parkinson’s disease (Camargo et al., 2015), multiple sclerosis (Marshall et al., 2014), and traumatic brain injury (Churchill et al., 2020; Costa et al., 2016; Mutch et al., 2016), among others. Vascular smooth muscle cell dysfunction attributed to vascular and neurodegenerative pathologies may also be identified by changes in CVR (Hayes et al., 2022; Khalil, 2010).

Non-invasive magnetic resonance imaging (MRI) has emerged as a promising tool for characterising anatomical and haemodynamic changes in the brain. In particular, functional magnetic resonance imaging (fMRI) based on blood oxygen level-dependent (BOLD) contrast acquired during a breath-hold (BH) task is a robust method to derive CVR maps in response to vasoactive stimuli (Pinto et al., 2021; Stickland et al., 2022; Zvolanek et al., 2023). The BH task leads to an elevation of the partial pressure of *CO*_2_ (*PaCO*_2_) in the blood, which triggers the dilation of blood vessels and leads to an increase in CBF, that can be measured with BOLD fMRI. The *PaCO*_2_ at the end of each exhalation (end-tidal *CO*_2_, *P*_*ET*_ *CO*_2_) is commonly used as a non-invasive surrogate of arterial blood *CO*_2_ (McSwain et al., 2010; Solis-Barquero et al., 2021; Takano et al., 2003). Consequently, it becomes possible to determine CVR by quantifying the ratio between the change in CBF and change in *P*_*ET*_ *CO*_2_, as depicted in Eq. (1).

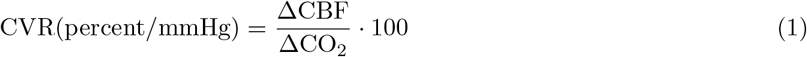

The CVR metric defined in Eq. (1), assumes that the CVR response occurs at the same time across the different brain regions. However, there are known differences in CVR delays across the brain, due to a variety of factors including different blood arrival times and tissue reactivity to *PaCO*_2_ along the vascular tree (Magon et al., 2009). These temporal features need to be taken into account to accurately characterise CVR amplitude across the brain (Pinto et al., 2016).

The most common way to model CVR amplitude and delay is through the application of time-shifted regressors in a general linear model (GLM) approach (Moia et al., 2020; Sleight et al., 2021). In this work, we compare this method with a novel variational Bayesian (VB) approach for CVR mapping which allows for the incorporation of prior information and simultaneous fitting of CVR amplitude and delay (Chappell et al., 2009). In this study, we evaluate the performance of these two methods on multi-echo BOLD fMRI data, acquired in ten subjects performing a BH task during ten repeated sessions each (Moia et al., 2021), to obtain subject-specific CVR and haemodynamic delay estimates, and their corresponding reproducibility metrics.

### 1.1 Lagged General Linear Model

The conventional approach to derive CVR from the BOLD fMRI time series is to use a general linear model (GLM), in which the acquired signal is separated into pertinent parameters of interest. In the GLM approach, the measured BOLD fMRI signal from each voxel, *Y*, is expressed as the sum of experimental design variables in a scaled design matrix comprised of regressors of interest (in this case the corresponding *P*_*ET*_ *CO*_2_ time course), and a combination of nuisance regressors, including the motion parameters and their temporal derivatives (denoted Mot), Legendre polynomials of up to the fourth order (denoted Poly), and rejected independent components (denoted *IC*_*rej*_) that have been orthogonalised with respect to the *P*_*ET*_ *CO*_2_ trace and the accepted components (denoted *IC*_*acc*_), and residual random noise (denoted *E*) as show in Eq. (2).

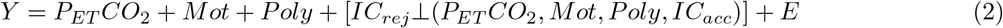

GLM is a univariate approach by which each voxel is treated as independent from one another, and it is assumed that the errors are random and follow a Gaussian distribution with a mean of zero.

To achieve accurate estimates of CVR amplitude using a GLM, it is important to also consider spatially variable haemodynamic delays (lags) between the *P*_*ET*_ *CO*_2_ regressor and the measured BOLD signal. To account for this, several GLMs are performed, each using a time-shifted *P*_*ET*_ *CO*_2_ regressor, and a single delay is selected for each voxel by selecting the shift that yielded the highest CVR amplitude (Moia et al., 2021; Stickland et al., 2021; Zvolanek et al., 2023).

### 1.2 Variational Bayesian Modelling

In Bayesian modelling, variables are treated as probability distributions as opposed to fixed values. Based on the Bayes theorem, the series of measurements, *y*, are used to determine the parameters, *w*, using the chosen model, *M*. The posterior probability of the parameters given the data and the model, i.e. *P* (*w*|*y, M*), is then a product of the likelihood of the data given *M* and *w*, i.e., *P* (*y*|*w, M*), the prior probability of the parameters given the model, i.e., *P* (*w*|*M*), and the evidence for the measurements in the chosen model, i.e. *P* (*y*|*M*), as shown in Eq. (3).

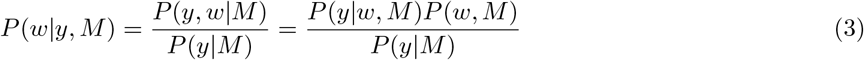

In most cases, it is not feasible to evaluate the posterior probability distribution analytically. Alternatively, a variational Bayesian (VB) approach implements this solution as derived by Chappell et al., 2009. In this case, an approximate posterior distribution *q*(*w*) can be computed and parameterized to define the form of the distribution on the parameters of interest, *w*. Then, the true posterior distribution can be inferred by minimising an error measure that quantifies the deviation of the approximate distribution *q*(*w*) from the exact Bayes’ posterior. The fit of the approximate distribution to the true distribution can be calculated using the variational free energy, *F* (which encodes the divergence between the approximate and true probability distributions) as shown in Eq. (4).

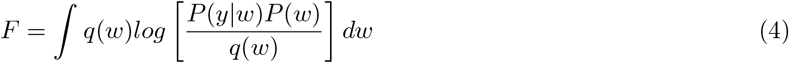

The correct estimation of *P* (*w*|*y*) is achieved by maximising free energy over *q*(*w*), which is equivalent to minimising the statistical dissimilarity, between *q*(*w*) and the true posterior distribution (Attias, 1999).

A common form for the prior distributions is a Gaussian distribution with a defined mean and variance as shown in Eq. (5) and Eq. (6). *P* (*θ*) has a normal distribution with mean *m*_*θ*_ and variance 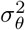 for the parameter of interest, *θ*, and *P* (*t*) is the temporal prior with mean *m*_*t*_ and variance 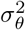.

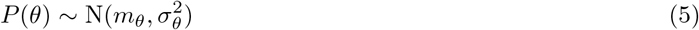

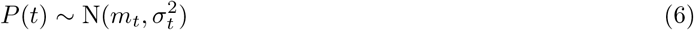

The mean and variance of the posterior distribution are commonly estimated using the priors and the data. In the context of deriving CVR maps from BOLD data, the VB approach replaces multiple GLMs with time-shifted regressors (lGLM) to a single model fit, where CVR delay is a parameter obtained simultaneously with CVR amplitude. Notably, VB inference also provides useful regularisation to deal with noisy imaging data and is far less computationally intensive than other Bayesian methods (Chappell et al., 2009).

## 2 Materials and Methods

### 2.1 Data Collection

Multi-echo BOLD fMRI data was previously acquired in ten subjects with no history of psychiatric or neurological disorders (5 females, range of 24-40 years of age). Each subject performed a BH protocol which was repeated for ten sessions over ten weeks (one each week), on a 3T Siemens PrismaFit scanner with a 64-channel head coil (340 scans, TR=1.5 s, TEs=10.6/28.69/46.78/64.87/82.96 ms, FA=70°, multiband=4, GRAPPA=2 with gradient echo reference scan, 52 slices with interleaved acquisition, Partial Fourier=6/8, voxel size=2.4×2.4×3 mm3) (Moia et al., 2021). The protocol consisted of eight repetitions of a BH task composed of four paced breaths of 6 s each, a 20 s BH, an exhalation of 3 s, followed by a recovery period of 11 s without pacing. Subjects were instructed prior to scanning about the importance of the exhalations preceding and following the apnoea to accurately characterise the *P*_*ET*_ *CO*_2_ (Pinto et al., 2021). During the fMRI acquisitions, exhaled *CO*_2_ and *O*_2_ levels were measured using a nasal cannula (Intersurgical) with an ADInstruments ML206 gas analyser unit. A complete detailed description of the participants and acquisition protocol are described in Moia et al., 2021.

### Data Preprocessing

The end-tidal *CO*_2_ peaks across the *CO*_2_ time courses were automatically and manually identified using custom scripts in Python 3.6.7 (Markello & DuPre, 2020; Moia et al., 2021) after high-pass filtering and downsampling. Linear interpolation between the end-tidal peaks was used and a cross-correlation bulk shift was applied to the time series to match *P*_*ET*_ *CO*_2_ values with the optimally-combined BOLD images. The *P*_*ET*_ *CO*_2_ signal was not convolved with a haemodynamic response function as the time-scale of the effect is assumed to be negligible during the gradual increase in *PaCO*_2_ that occurs throughout a BH.

Preprocessing of the MRI data was done in AFNI (Cox, 1996), FSL (Jenkinson et al., 2012), and ANTs (Tustison et al., 2014) using custom scripts (Moia, 2022; Moia et al., 2021). The T2-weighted images were skull-stripped and the obtained brain map was coregistered to the T1w space. The latter was tissue-segmented in grey matter (GM), white matter (WM), and cerebral spinal fluid (CSF), and normalised to the MNI template (FSL 2mm). The first 10 volumes of the MRI data were discarded to remove the rest portion of the task data, and ensure that a steady state of magnetisation was achieved. After motion realignment, the multiple echo time series were optimally combined voxelwise by weighting each time series by its T2 value, obtaining a single T2^*^w timeseries. Multi-echo independent component analysis decomposition was performed using tedana (DuPre et al., 2019), and manual classification of the independent components was conducted by experts based on temporal, spatial, and spectral features. Contrarily to Moia et al., 2021, voxelwise denoising was applied via nuisance regression before subsequent data analysis. The set of nuisance regressors included: the motion realignment parameters and their temporal derivatives, Legendre polynomials up to 4th order, and the rejected independent components from ME-ICA which were previously orthogonalized with the accepted independent components and the *P*_*ET*_ *CO*_2_ signal at delay time zero. This approach was chosen so that both the VB and lGLM could be compared with the same denoising and preprocessing steps. It should be noted that simultaneous fitting of the nuisance regressors and the regressor of interest is preferable, but is expected to be very similar to the sequential modelling approach (Lindquist et al., 2019; Moia et al., 2020).

### 2.3 Data Analysis

Data analysis was performed using Quantiphyse, an analysis and visualisation platform for medical imaging data (Quantiphyse, 2023). The CVR toolbox in Quantiphyse was used for the VB and lGLM analyses to estimate the CVR amplitude and delay maps for each subject and session. To account for measurement delay, Quantiphyse applied a bulk time-shift to individual *P*_*ET*_ *CO*_2_ traces, where at the time of the bulk shift corresponds to a delay of 0 s. This time-shift was obtained by selecting the shift that yielded the highest cross-correlation between *P*_*ET*_ *CO*_2_ trace and the corresponding average whole-brain fMRI signal. For the lGLM approach, we established a window for the candidate CVR delays of *±* 8 s from the bulk time-shift with a 0.25 s timestep. The VB analysis was done with the BOLD timeseries modelled as a scaled difference between the *P*_*ET*_ *CO*_2_ timeseries at a delay shift *t*_*delay*_ relative to the minimum *P*_*ET*_ *CO*_2_ with error *ϵ*, as shown in Eq. (7).

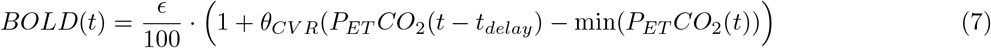

The VB analysis used 10 iterations, starting with wide non-informative priors. The prior distribution for the CVR was *P* (*θ*) ∼ *N* (1, 2000), and the prior for the delay was *P* (*t*) ∼ *N* (0, 100).

### 2.4 Statistical Analysis and Reproducibility

All CVR and delay maps were co-registered to the MNI152 template using ANTs tool and nearest neighbour interpolation (Grabner et al., 2006). A revised linear mixed effects (LMEr) model was then applied voxel-wise, taking into account variability within and across subjects (Chen et al., 2013). The LMEr model was formulated as presented in Eq. (8) where *V ar* represents the variable CVR or delay of each voxel.

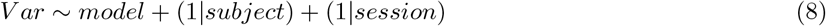

Z-scores produced by the LMEr calculation were thresholded to investigate statistical significance, defined as cluster-corrected p-value *<* 0.01.

Reproducibility of the CVR and delay maps for each method was assessed by computing the intraclass correlation coefficient (ICC) considering two-way random effects, absolute agreement, and a single measurement (ICC(2,1)) with AFNI’s tool 3dICC (Chen et al., 2018; Koo & Li, 2016). The ICC(2,1) is calculated as shown in Eq. (9) where *MS*_*sub*_ is the mean square for subjects, *MS*_*ses*_ is the mean square for sessions, *MS*_*n*_ is the mean square for residuals, *k* is the number of sessions, and n is the number of subjects (Shrout & Fleiss, 1979).

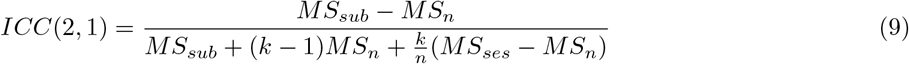

Note that a high ICC score (up to 1) indicates high reproducibility, where intrasubject variability is lower than intrasession variability. For fMRI studies, an ICC score below 0.40 is considered poor, 0.40-0.59 is considered fair, above 0.60-0.74 is good, and above 0.75 is excellent (Aron et al., 2006; Cicchetti, 2001; Eaton et al., 2008).

The ICC analysis was first conducted voxelwise across the lGLM and VB maps for both methods. A regional analysis was also performed using the MNI-maxprob-thr25 brain atlas at 2.5mm in FSL (Collins et al., 1995; Mazziotta et al., 2001), applied to the standardised CVR and delay maps. The ICC scores were calculated from the mean and median values within each region across all subjects and sessions.

## 3 Results

Four subjects were excluded due to incomplete *P*_*ET*_ *CO*_2_ traces in one or more of the 10 sessions. Excluded traces were most often the result of inadequate execution of the exhalations preceding and following the BH which prevented accurate determination of the *P*_*ET*_ *CO*_2_ traces.

### 3.1 Cerebrovascular Reactivity and Delay Maps

The CVR amplitude and delay maps derived for a representative subject and session are presented in Fig. 1 and Fig. 2, respectively, for each method. Figures showing the CVR and delay maps for the same subject across all sessions with both methods are available in Fig. S1 and Fig. S2 of the Supplementary Material.

**Figure 1:**
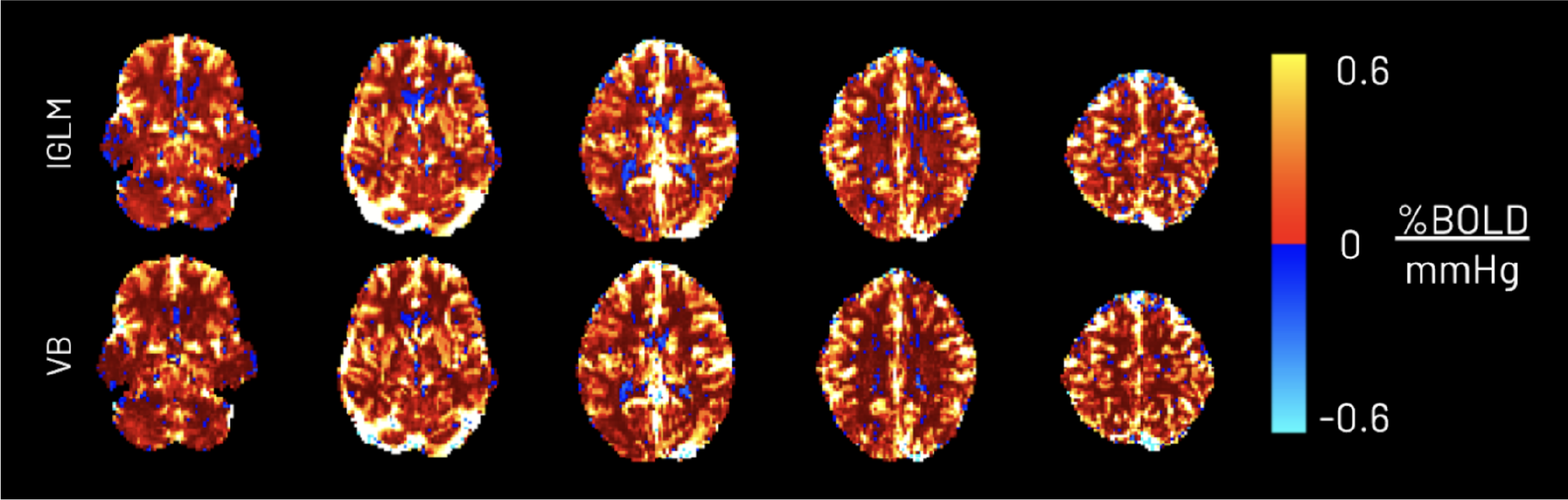
CVR amplitude maps obtained using the lagged-GLM (lGLM) and variational Bayesian (VB) analyses for a representative subject and session.

**Figure 2:**
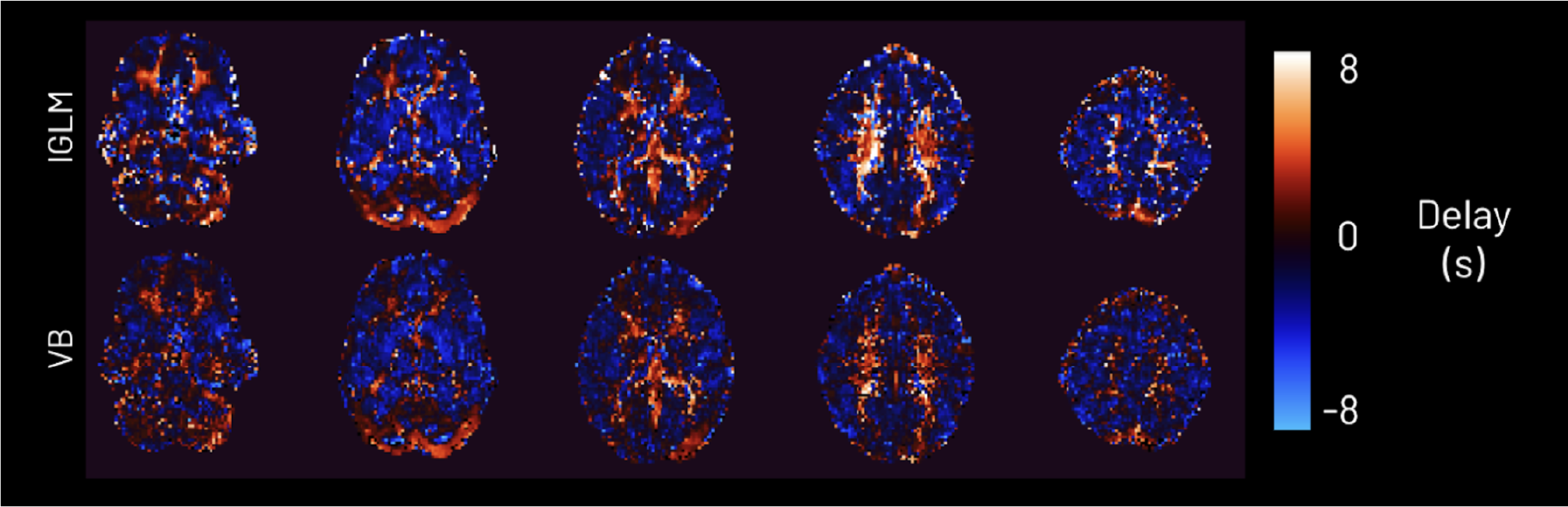
CVR haemodynamic delay maps obtained using the lagged-GLM (lGLM) and variational Bayesian (VB) analyses for a representative subject and session.

Scatter plots comparing the CVR values derived by both methods of a representative session are presented in Fig. 3 along with the best fit line corresponding to *y* = 0.93*x* − 0.0002 and a Pearson R of 0.98. The same comparison for the delay maps are presented in the top plot of Fig. 4. The lGLM delay map calculation was repeated at a higher time resolution (with a time step of 0.025 s from -8 s to 8 s) for the representative case and the scatter plot comparison is presented in the bottom plot of Fig. 4. The best fit line for both delay value comparisons corresponds to the equation *y* = 0.36*x* − 0.35 and a Pearson R of 0.34. The least sum of absolutes fit for the CVR, and both delay maps correspond to Spearman rho constants of 0.94 and 0.65 respectively.

**Figure 3:**
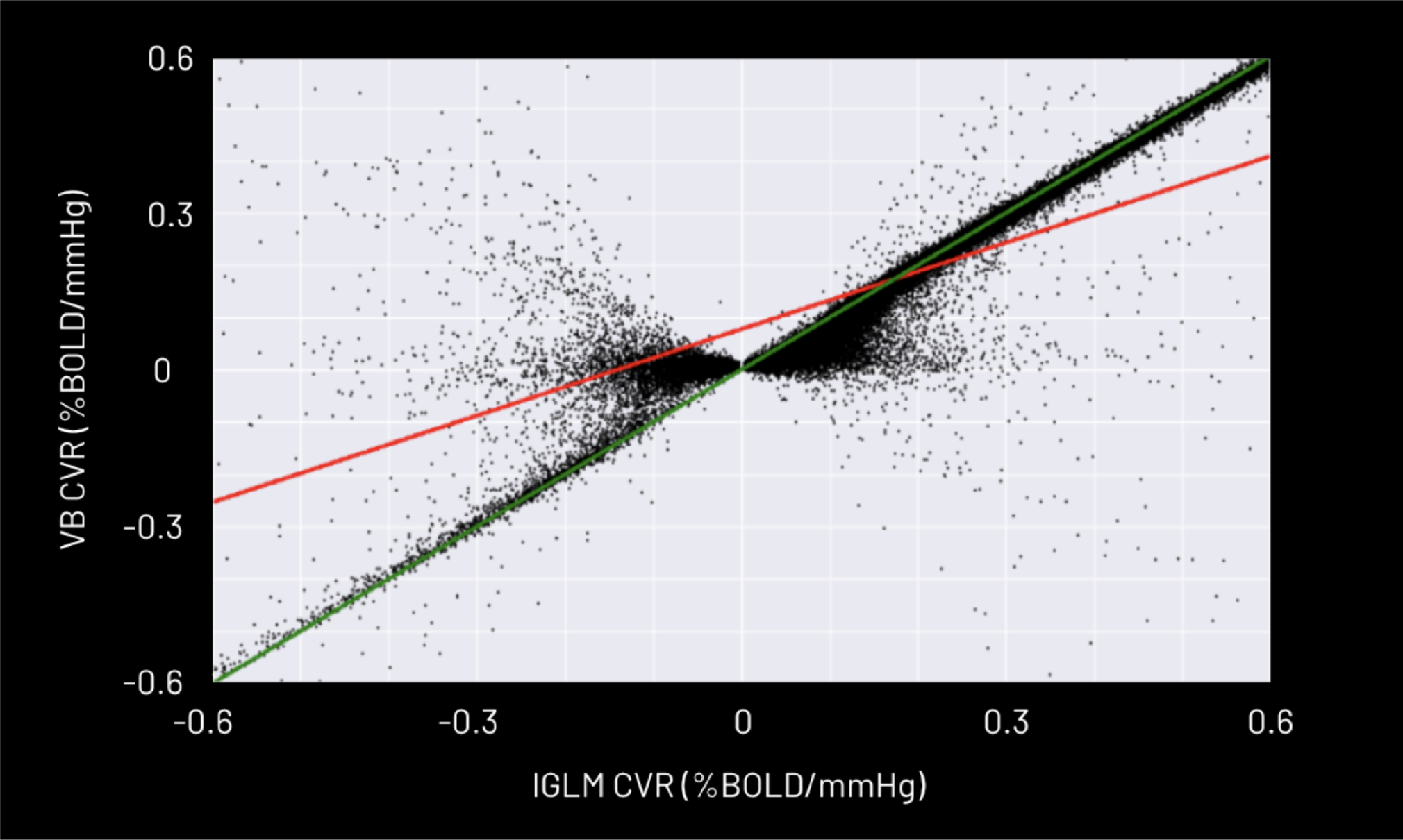
Scatter plot of CVR amplitude values obtained using the variational Bayesian (VB) method as a function of those obtained using the lagged general linear model (lGLM) approach for a representative subject and session. The least squares fit line is plotted in red which corresponds to the equation *y* = 0.93*x* − 0.0002 and a Pearson R of 0.98. The least sum of absolutes fit of this data corresponds to a Spearman rho constant of 0.94. The y=x line is plotted in green.

**Figure 4:**
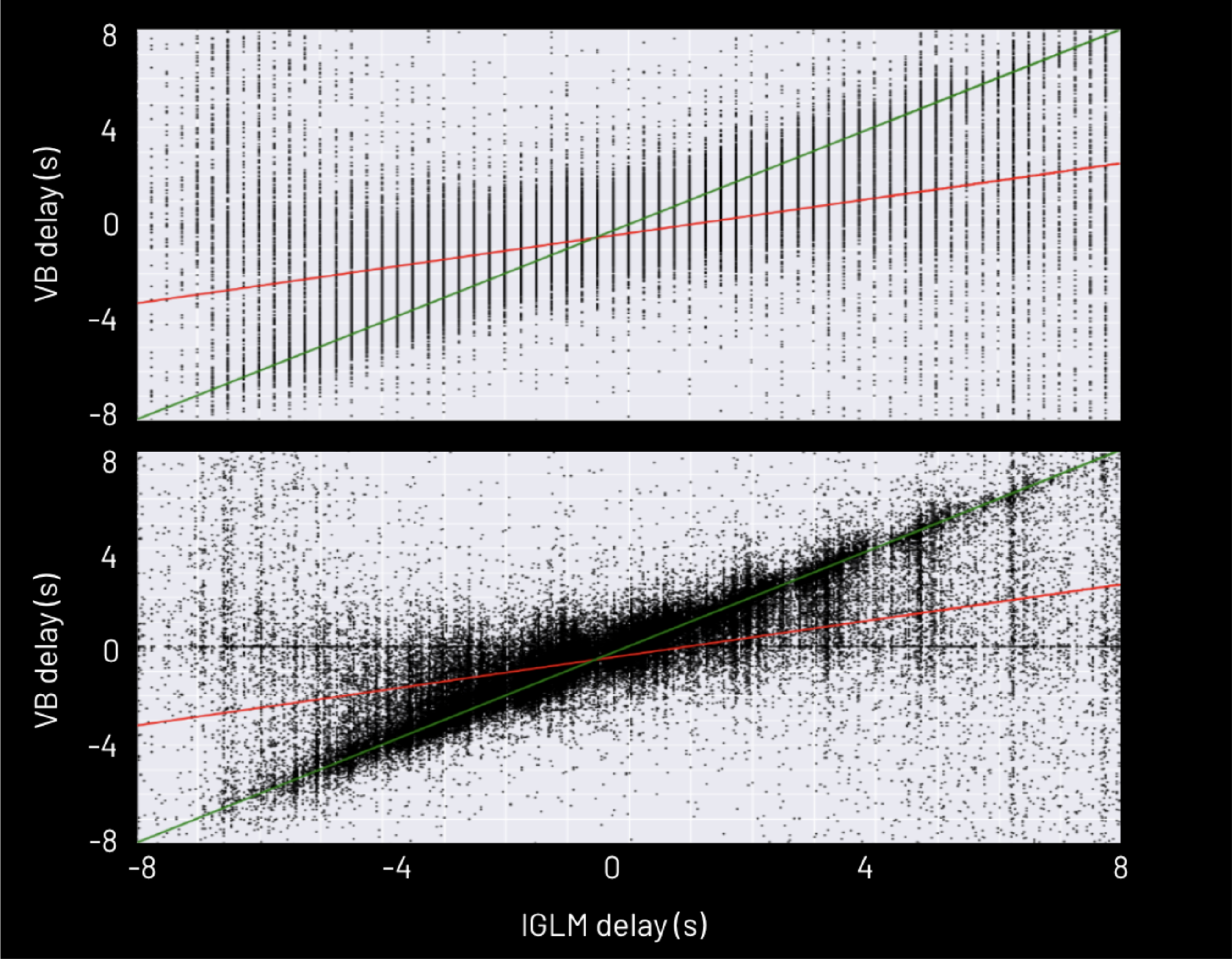
Scatter plots of haemodynamic delay values obtained using the variational Bayesian (VB) method as a function of those obtained using the lagged-GLM (lGLM) approach for a representative subject and session. The scatter plots show the lGLM delay values calculated using a 0.25s timestep (top) and a 0.025s timestep(bottom). The least squares fit line is plotted in red which corresponds to the equation *y* = 0.36*x* − 0.35 and a Pearson R of 0.34 for both timesteps. The least sum of absolute fits of these comparisons both correspond to a Spearman rho constant of 0.65.

### 3.2 Statistical Analysis

The results of the revised linear mixed effects (LMEr) analysis are presented in Fig. 5. It should be noted that the range of -0.03 to 0.03 percent/mmHg is much smaller than the range of CVR values (-0.6 to 0.6 percent/mmHg). Notably, there are no statistically significant differences between the CVR values of the lGLM and VB methods in grey matter (using *p <* 0.01, FDR corrected). Statistically significant differences between the methods (*p <* 0.01, FDR corrected) are apparent in white matter and CSF, especially around the ventricles. P-values were adjusted for control of false discovery rate (FDR) (Benjamini & Hochberg, 1995), and then compared against an alpha of 0.01 to determine significance. The thresholded z-score maps for the CVR and delay LMEr comparison are available in Fig. S3 of the Supplemental Material.

**Figure 5:**
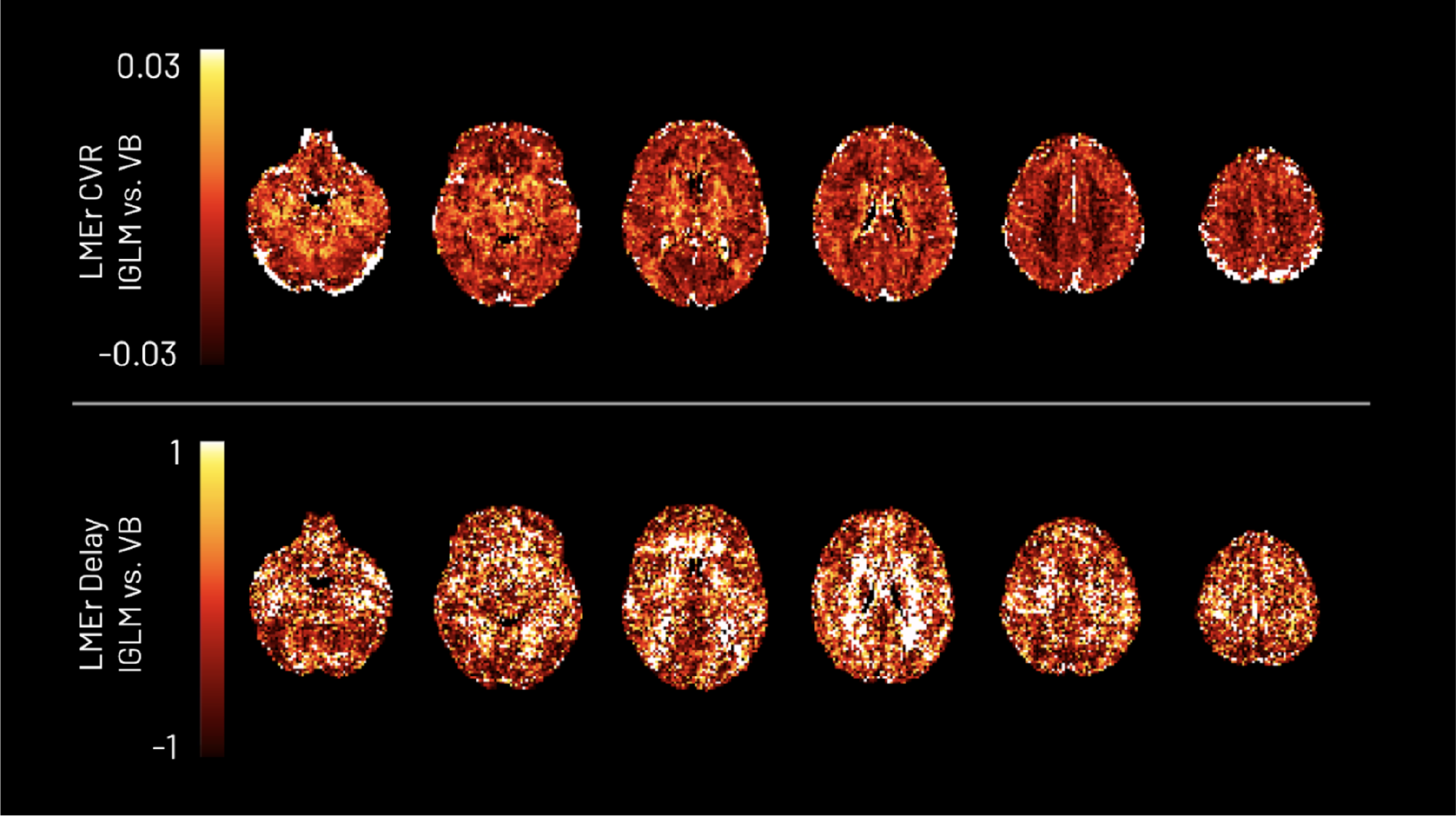
Revised linear mixed-effects (LMEr) pairwise comparison between the lagged-GLM (lGLM) and variational Bayesian (VB) analyses. It should be noted that the range of -0.03 to 0.03 is much smaller than the range of CVR values (-0.6 to 0.6 percent/mmHg) and delay values (-8 to 8 s).

### 3.3 Reproducibility

#### 3.3.1 Voxelwise Reproducibility

The ICC scores in the CVR amplitude maps are presented in Fig. 6 and indicate a high reproducibility with higher intersubject variability than intrasubject variability in both methods. Poor reproducibility (*<* 0.4) is illustrated for the delay maps for both methods, attributed to high variability in fitting this parameter and differences in bulk shifts between sessions.

**Figure 6:**
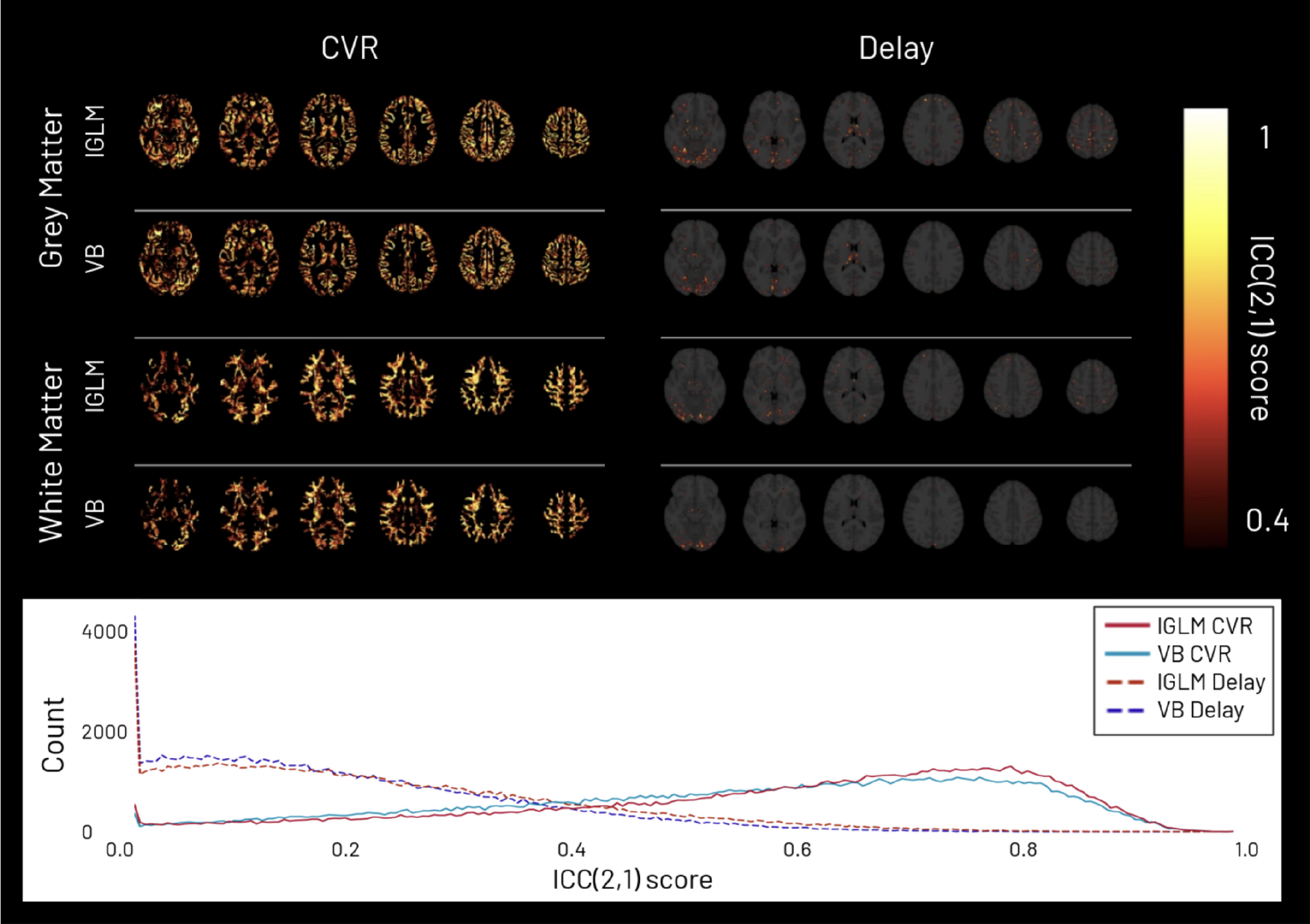
Intraclass correlation coefficient ICC(2,1) maps of CVR amplitude (left) and delay maps (right) for the lagged-GLM (lGLM) and variational Bayesian (VB) methods in grey matter and white matter. The maps are thresholded at 0.4 since scores lower than it indicate poor reproducibility. The bottom row depicts the whole brain distributions of ICC scores across voxels for CVR amplitude (solid lines) and delay (dashed lines) for the lGLM (red) and VB (blue) methods.

#### 3.3.2 Regional Reproducibility

The regional ICC scores for both methods are presented in Table 1 using the mean and median values within each region. Notably, a handful of outliers using the lGLM method skewed the mean value in some regions, significantly reducing the ICC score (denoted by ^*^ in Table 1). The median was used to remove the effect of outliers. The group-level mean and median CVR and delay values for both methods in each region are available in Table S1 of the Supplemental Material.

**Table 1:**
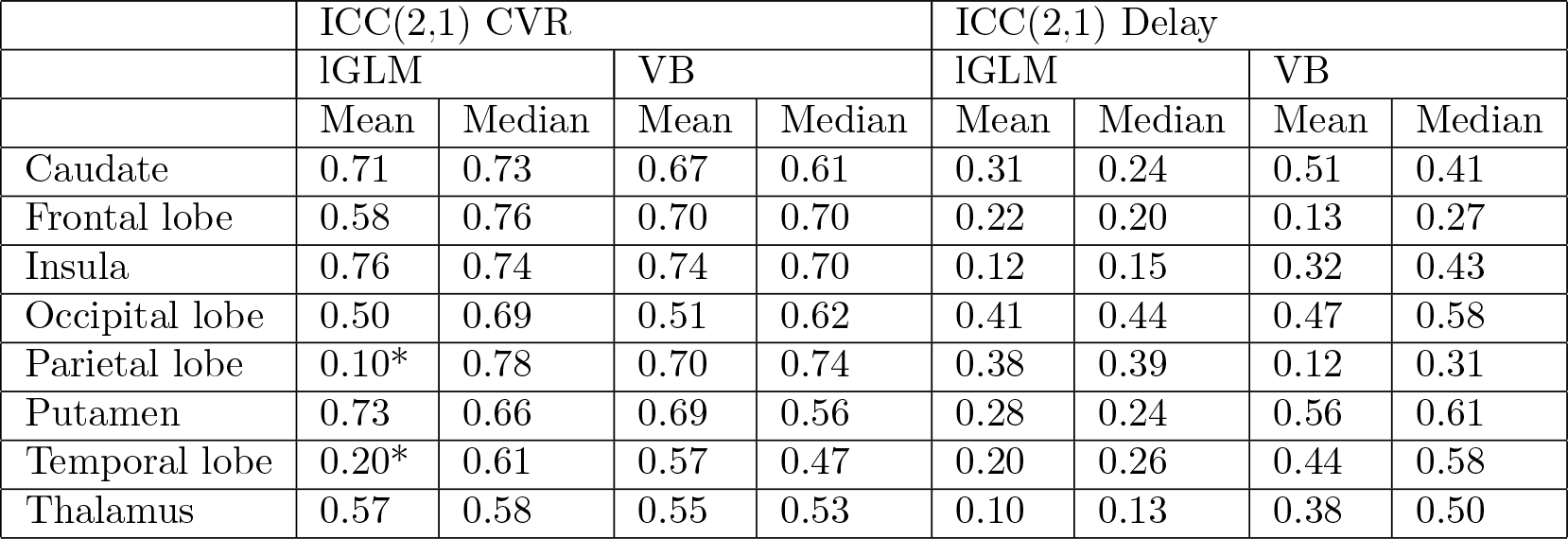
Regional intraclass correlation coefficient ICC(2,1) values of the CVR amplitude and delay maps for lGLM and VB methods in 8 MNI-atlas regions of the brain. ^*^ denotes regions containing significant outlier values in the mean calculation.

## 4 Discussion

In this study, we compared two modelling approaches, the lagged-GLM (lGLM) and variational Bayes (VB), to estimate CVR amplitude and delay maps based on BOLD fMRI data acquired during a BH protocol. To the authors’ knowledge, this is the first time these approaches have been compared in the context of CVR and delay mapping using multi-echo BOLD fMRI.

Regarding our BOLD-fMRI data preprocessing, multi-echo independent-component analysis denoising was used to remove components exhibiting noise-like signal decay patterns across echos (DuPre et al., 2019). An optimal combination of the echos was used as it has been shown to improve BH-induced CVR mapping sensitivity, specificity, repeatability, and reliability (Cohen & Wang, 2019). For both modelling approaches, a conservative nuisance regression approach was used prior to the CVR fitting to preserve the BOLD variance associated with local CVR responses while still adequately removing noise-related effects from the time series (Moia et al., 2021).

The main difference between the lGLM and VB modelling strategies is that the VB method treats the inputs and outputs as probability distributions, and simultaneously fits the CVR amplitude and temporal delay parameters. Visually comparing the maps derived using the lGLM and VB methods indicates that the output CVR maps are very similar between methods. The LMEr analysis confirms that the CVR amplitude maps do not show statistically significant differences between methods in grey matter (using *p <* 0.01, FDR corrected). Within these grey matter regions, the CVR values are highly comparable to results from previous studies that also take regional haemodynamic delays into account (Lipp et al., 2015; Moia et al., 2021; Pinto et al., 2016). White matter and CSF show statistically significant differences between methods (*p <* 0.01, FDR corrected), likely illustrating that the VB and lGLM methods treat low SNR voxels differently. This is also seen in the scatter plot comparison where low signal voxels result in low CVR amplitude values in the lGLM method, but are close or equal to zero in the VB method.

The LMEr of the delay maps shows a variation of *±*1 s of the mean whole-brain delay time between methods. The scatter plot comparison of the delay maps showcases how the discrete time step in lGLM approach forces all estimates into bins as opposed to allowing for a continuous distribution. At the expense of computational resources, this was improved by increasing the time step in one representative subject. Notably, in comparing the delay values for both methods, the VB method forces more voxels towards a delay of 0 s, attributed to the prior distribution reverting low SNR voxels to the prior distributions. Consistent with previous studies, we find considerable variations in the delay across the brain with both methods, supporting the use of modelling strategies that include temporal aspects of the CVR response (Andrade et al., 2006; Bright et al., 2009; Leoni et al., 2008).

The voxelwise and regional CVR ICC scores in this analysis are similar to previous work and illustrate high reproducibility for both methods (Bright & Murphy, 2013; Magon et al., 2009; Moia et al., 2021; Pinto et al., 2016). The regional CVR amplitude ICC values for the VB method (Mean: 0.51-0.74; Median: 0.47-0.74) are all above 0.4, considered the acceptable minimum in fMRI studies (Aron et al., 2006; Cicchetti, 2001; Eaton et al., 2008). The regional ICC values for the lGLM method (Mean: 0.10-0.76; Median: 0.58-0.78) highlighted large outliers that skewed the mean and resulting ICC scores in the parietal and temporal lobes, while all other regions are above 0.4. This issue was overcome by using the median CVR values within each region, which brought all ICC values above 0.4 (fair) and most above 0.6 (good). The ICC values are comparable to previously reported regional CVR ICC, notably showing higher reproducibility in the frontal and parietal lobes, and lower reproducibility in the parietal and temporal lobes (Bright and Murphy, 2013).

The voxelwise delay map ICC scores are lower than previously reported (Moia et al., 2021). Fewer subjects were included in our analysis compared to Moia et al., 2021, which is expected to have altered the results of the reproducibility assessment. Furthermore, we are using a different ICC test model since in Moia et al., 2021, the t-statistic maps associated with the estimation of the CVR using the lGLM method were attributed to the delay values and were included in their ICC calculation (Chen et al., 2017; Moia et al., 2021). The t-statistics were excluded from the reproducibility comparison for fairness to both methods since t-statistic maps are not presently calculated during the VB implementation in Quantiphyse. Without the t-statistic, both the lGLM and VB methods show good reproducibility in CVR maps, but poor reproducibility in the delay maps, likely due to high-variance estimates between sessions from the ICC calculation itself as a result of session-specific temporal shifts.

The regional delay ICC results for the VB method (Mean: 0.12-0.56; Median: 0.27-0.61) and lGLM method (Mean: 0.10-0.41; Median: 0.13-0.44) range from fair (ICC *>* 0.4) to poor ICC levels (ICC *<* 0.4). Notably, the regional delay reproducibility is higher than the voxelwise reproducibility as it averages over noisy voxels. Fair delay reproducibility was found in the caudate, insula, occipital lobe, putamen, temporal lobe, and thalamus when using the median VB-derived values. Only the occipital lobe showed fair reproducibility when using the lGLM-derived delay values.

Both methods reported primarily positive CVR amplitude values in GM; however, negative CVR values are visible in WM and CSF. This negative CVR might originate from inaccurate modelling in noisy voxels with reduced CVR-related changes and/or other flow and volume changes (e.g., CSF). The ‘vascular steal’ effect has also been reported as an underlying physiological cause of negative CVR, which may be used as a biomarker of pathology (Poublanc et al., 2013). However, vascular steal does not appear to be an expected physiological event in the absence of pathology (Arteaga et al., 2014), and due to the consistent pattern of negative CVR values in the WM and CSF throughout all healthy subjects, the negative CVR voxels are not probably caused by this effect. However, the mechanisms behind negative CVR in specific brain tissues in healthy subjects are not yet explained; hence, further work is still required to validate this observation. Practically speaking, the VB method can be more time efficient at scale as it fits the CVR and delay simultaneously, while the lGLM approach requires several GLMs to assess both parameters. Additionally, the lGLM approach uses a step-fit and requires the user to select the delay range and step size a priori. To illustrate this, the analysis of a subset of the data was timed on a shared high-performance computer with an Intel Xeon and a CPU with 32 cores and 132 GB of RAM. For 10 sessions, the VB method took 69 mins and the lGLM method took 130 mins, marking over an 80% increase in performance of the VB method compared to the lGLM method. We expect the computation cost to scale with the number of sessions because no parallelization was implemented. To approach the continuous distribution of delay values generated by the VB modelling, the lGLM method will take exponentially more time as the model recomputes a GLM for each voxel at each specified delay value in a discreet manner.

When using Quantiphyse, the denoising and voxelwise response estimation could not take place simultaneously since multiple regressors are not yet implemented. Simultaneous regression has previously been shown to improve the output accuracy as it ensures that the degrees of freedom and the interactions between regressors are accounted for (Jo et al., 2013; Lindquist et al., 2019). This is a difference from Moia et al., 2020 and Moia et al., 2021 which showed that simultaneous modelling is more appropriate to account for potential correlations between modelling regressors in BOLD fMRI. Future implementations of Quantiphyse could address this by allowing for multiple regressors.

For this study, we used wide non-informative priors that ultimately resulted in parameter estimates that are very similar to a least squares fit, with an additional time dimension (Syversveen, 1998). However, an advantage of the Bayesian approach is that prior information can be included to constrain the model to physiologically relevant results, and different models can be defined for the VB method to fit (Bolstad, 2009). Future work intends to explore different models and prior distributions in addition to priors for a noise model to deal with complex noise patterns that are not constant across the measurements (Chappell et al., 2009). This may be particularly useful for analysing task-based functional MRI data, where changes in stimuli and tasks may introduce different levels of noise throughout an acquisition. In the case of noisy data, variance maps can be calculated during the VB method and may be useful to assess the confidence in the VB model fit, however minimal trends were found to distinguish between tissue types or SNR in the CVR or delay maps and therefore were not used in this analysis (see Fig. S4 of the Supplemental Material). More research into confidence metrics on both the VB and lGLM methods is warranted as it would benefit their refinement and clinical uses.

As in the case of both VB and lGLM approaches, linear models are the most common form for generating CVR maps (Sleight et al., 2021). However, CVR has a sigmoidal non-linear relation to the *P*_*ET*_ *CO*_2_, and BH-induced hypercapnia can also have a complex shape in terms of temporal and amplitude responses through the brain due to multiple physiological factors (Bhogal et al., 2014; Magon et al., 2009). Accounting for these non-linear components is an important next step to improve the full characterisation of the CVR response, and demonstrates a potential benefit of the VB approach which can be expanded to allow for non-linear modelling.

## 5 Conclusion

Both lGLM and VB modelling strategies provide robust CVR amplitude and delay maps for breath-hold BOLD fMRI data, with no statistically significant differences observed between them in grey matter. Noteworthy distinctions are apparent for low CVR amplitude, attributed to how each method addressed noisy voxels, particularly in white matter and cerebral spinal fluid. The CVR mapping was found to be highly reproducible for both methods. Despite higher variability in haemodynamic delay maps between sessions for both methods, the VB method exhibited fair reproducibility in more regions than the lGLM method. Furthermore, the VB method estimated a continuous delay map, which is discrete for the lGLM method, and with a delay timestep of 0.25 s, the VB estimation performed 80% faster than the lGLM. More research is still required to identify goodness of fit and confidence metrics for both approaches, as well as to discriminate against low SNR voxels. Future work will include implementing different physiologically relevant priors and incorporating non-linear modelling into Bayesian CVR modelling. This study underscores the utility of variational Bayesian modeling for CVR mapping, emphasizing its potential to interpret BOLD fMRI dynamics in cerebrovascular health studies and clinical applications.

## 6 Data and Code Availability

The methods pipeline for this analysis is available at https://github.com/genhayes/vbayes_lglm_cvr_pipeline, and all of the MRI images, physiological data, and manual classification used in this study are available on OpenNeuro (Moia et al., 2020). The original preprocessing and analysis scripts are available at https://github. com/smoia/EuskalIBUR dataproc. Additional tutorials explaining how to use Quantiphyse are available at https://github.com/physimals/quantiphyse.

## 7 Author Contributions

**Genevieve Hayes**: Conceptualization, Methodology, Software, Formal Analysis, Writing – Original Draft, Writing – Revising and Editing, Visualization, Funding Acquisition. **Daniel P. Bulte**: Methodology, Resources, Investigation, Writing – Revising and Editing. **Stefano Moia**: Data Curation, Software, Formal Analysis, Investigation, Writing – Revising and Editing, Visualisation. **Martin Craig**: Software, Methodology, Resources, Writing – Revising and Editing. **Michael Chappell**: Software, Methodology, Resources, Writing – Revising and Editing. **Eneko Uruñuela**: Software, Investigation, Data Curation, Writing – Revising and Editing. **Sierra Sparks**: Investigation, Writing – Revising and Editing. **César Caballero-Gaudes**: Conceptualization, Methodology, Data Curation, Resources, Investigation, Writing – Revising and Editing, Supervision, Project Administration, Funding Acquisition. **Joana Pinto**: Conceptualization, Methodology, Resources, Investigation, Writing – Revising and Editing, Supervision, Project Administration, Funding Acquisition.

## 9 Acknowledgements

This research was funded by the Clarendon Fund, the Engineering and Physical Sciences Research Council UK (EP/S021507/1 and EP/P012361/2) the Spanish Ministry of Economy and Competitiveness (RYC-2017-415 21845), the Basque Government (BERC 2018-2021, PIB 2019 104), the Spanish Ministry of Science, Innovation and Universities (PID2019-105520GB-100), the GliMR 2.0 COST Action (CA18206), the European Union’s Horizon 2020 research and innovation program (Marie Sk-lodowska-Curie grant agreement No.713673), and a fellowship from La Caixa Foundation (ID 100010434, fellowship code LCF/BQ/IN17/11620063). The authors would like to thank the Basque Centre on Cognition, Brain and Language for its support.

