## Supplemental Figures and Tables for "Bayesian Modelling Approaches for Breath-Hold Induced Cerebrovascular Reactivity"

### Supplemental Material

#### Cerebrovascular Reactivity and Delay Maps Across Sessions

All sessions for one representative subject are presented in Fig. S1 and Fig. S2, respectively, for both the variational Bayesian (VB) and lagged general linear model (lGLM) methods.

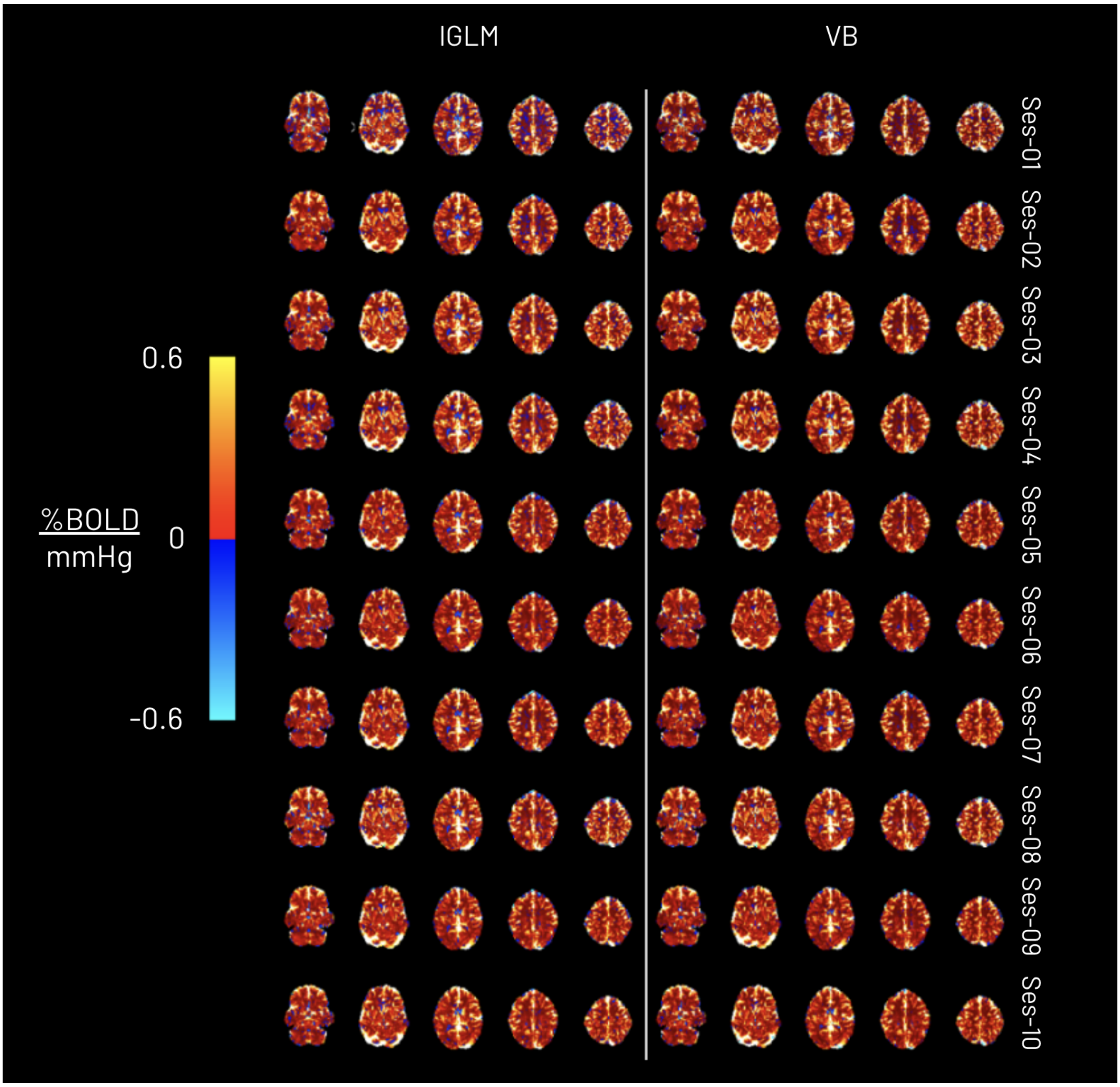


Figure S 1: CVR amplitude maps obtained using the lagged-GLM (lGLM) and variational Bayesian (VB) analyses for all the sessions of a representative subject (subject 002). Each row represents a session.

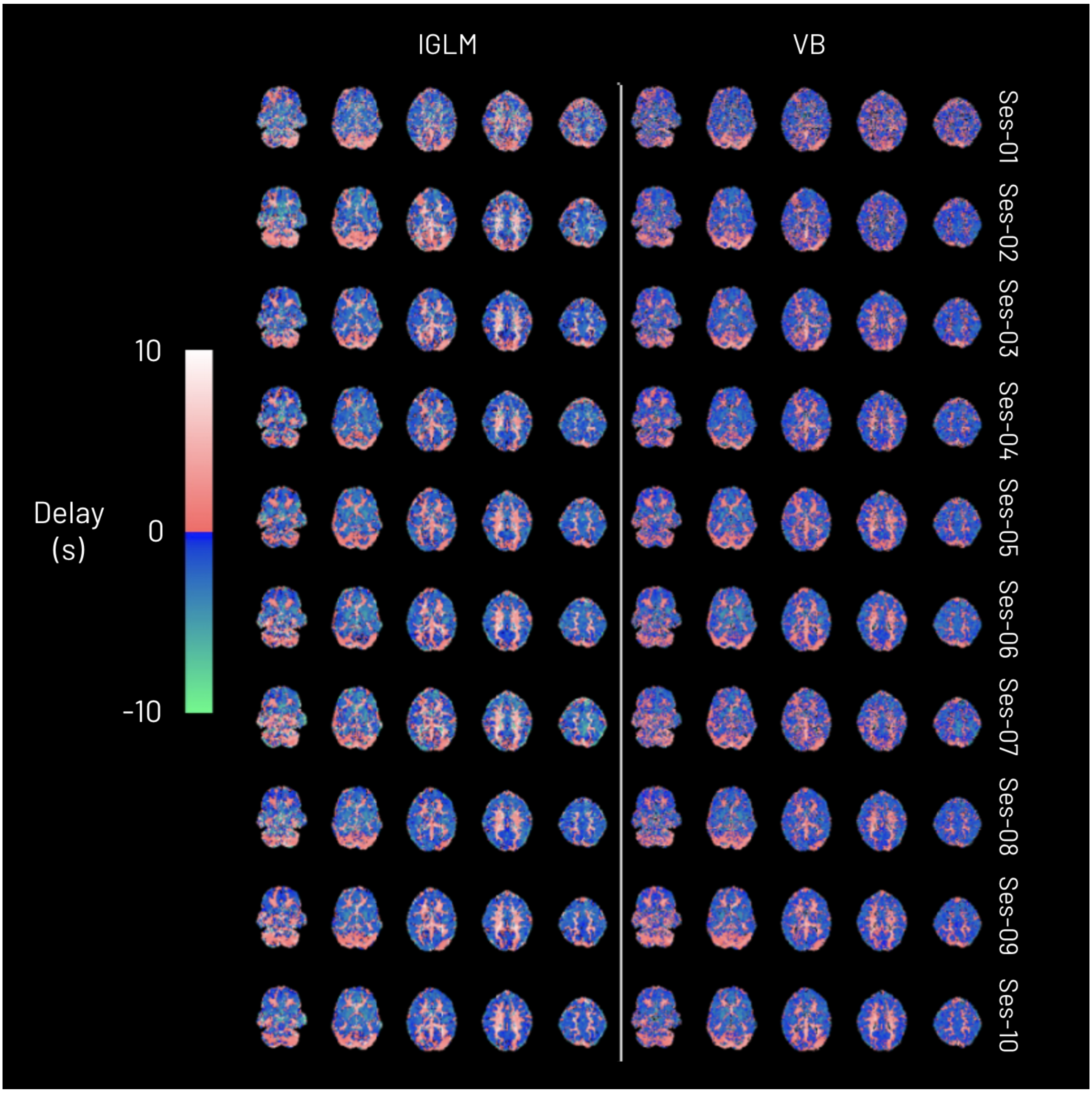


Figure S 2: CVR haemodynamic delay maps obtained using the lagged-GLM (lGLM) and variational Bayesian (VB) analyses for all the sessions of a representative subject (subject 002). Each row represents a session.

#### LMEr Z-scores

The thresholded z-score maps for the CVR and delay LMEr comparison are presented in Fig. S3. The absolute value of the z-score is thresholded at 2.63 for the CVR map and 2.60 for delay map, corresponding to a cluster-corrected p-value of 0.01.

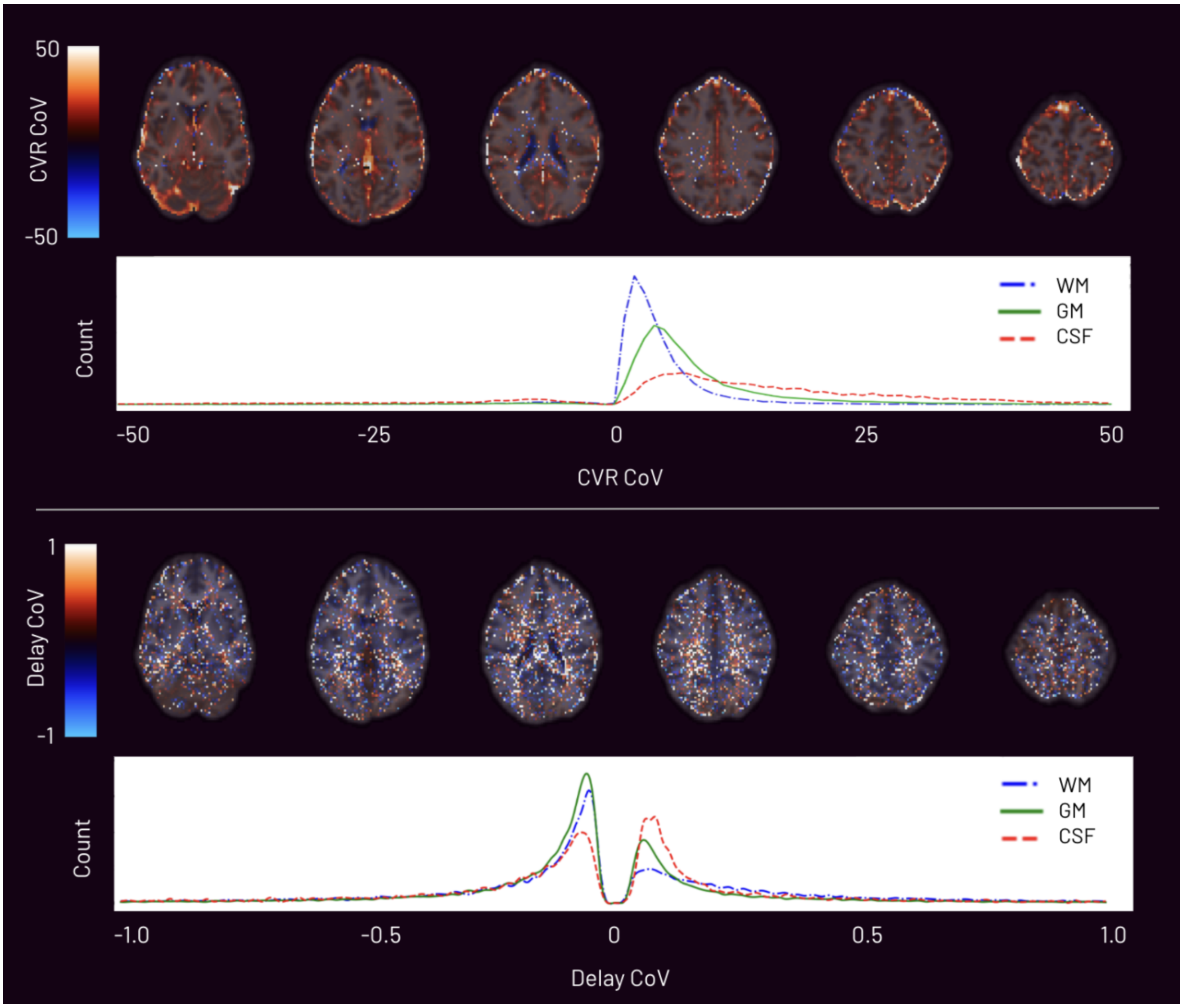


Figure S 3: CVR and delay z-score maps from the LMEr pairwise comparison between the lagged-GLM (lGLM) and variational Bayesian (VB) analyses. The absolute value of the z-score is thresholded at 2.63 for the CVR map and 2.60 for delay map, corresponding to a cluster-corrected p-value of 0.01.

#### Group-Level Mean and Median CVR and Delay Values

The CVR and delay values within each of the 8 regions of the MNI brain atlas (MNI-maxprob-thr25 brain atlas at 2.5mm in FSL) are shown in Table S1.

Table S 1: Regional CVR amplitude and delay values for lGLM and VB methods in 8 MNI-atlas regions of the brain.

| | CVR (%BOLD/mmHg) | | | | Delay values (s) | | | |
|---|---|---|---|---|---|---|---|---|
| | lGLM | | VB | | lGLM | | VB | |
| | Mean | Median | Mean | Median | Mean | Median | Mean | Median |
| Caudate | 0.104 | 0.127 | 0.094 | 0.103 | -0.128 | -1.104 | -0.110 | -0.524 |
| Cerebellum | 0.153 | 0.132 | 0.174 | 0.112 | 1.397 | 1.692 | 0.958 | 0.862 |
| Frontal lobe | 0.199 | 0.161 | 0.200 | 0.144 | -0.848 | -1.667 | -0.601 | -0.597 |
| Insula | 0.172 | 0.184 | 0.162 | 0.172 | -1.004 | -1.596 | -0.757 | -1.01 |
| Occipital lobe | 0.237 | 0.155 | 0.220 | 0.141 | -0.268 | -0.633 | -0.101 | -0.212 |
| Parietal lobe | 0.189 | 0.157 | 0.197 | 0.141 | -0.727 | -1.183 | -0.468 | -0.530 |
| Putamen | 0.143 | 0.152 | 0.129 | 0.133 | -1.483 | -2.254 | -0.989 | -1.276 |
| Temporal lobe | 0.170 | 0.120 | 0.175 | 0.096 | -0.280 | -0.667 | -0.153 | -0.299 |
| Thalamus | 0.191 | 0.152 | 0.177 | 0.132 | -0.396 | -1.229 | -0.276 | -0.631 |

#### Variance and Coefficients of Variation

The VB method computes variance maps for each model parameter, detailing an uncertainty metric in the posterior outputs. Since the variance scales with the amplitude of the parameter, a parameter known as the coefficient of variation (CoV) was calculated by dividing the variance by the parameter of interest as shown in Fig. S4 for the CVR and delay CoV.

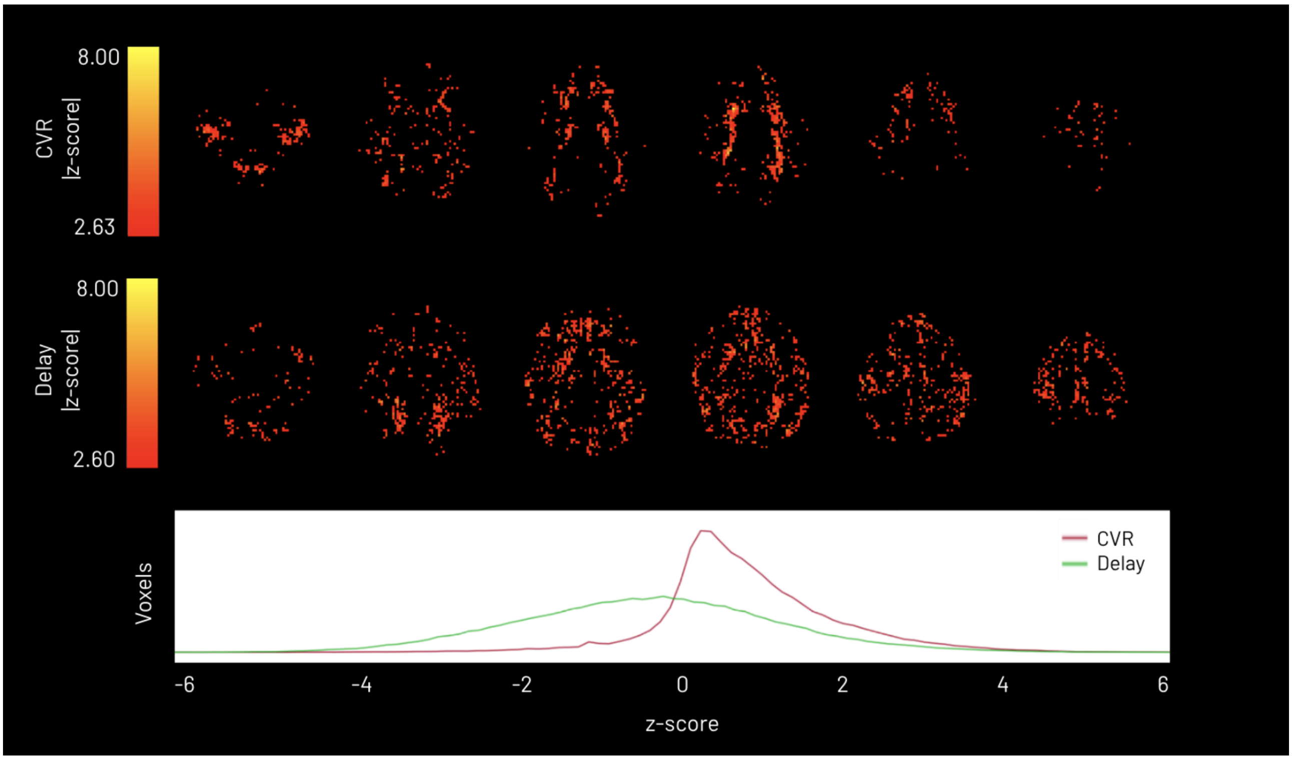


Figure S 4: Coefficient of variation (CoV) map of variational Bayesian-derived CVR (top) and delay (bottom) for a representative subject and session. The respective CoV histogram distributions are plotted of grey matter (green, solid), white matter (blue, dashed), and cerebral spinal fluid (red, long-dashed)
